# NPR1 protects young leaves from systemic salicylic acid-induced damage during bacterial infection

**DOI:** 10.1101/2023.08.21.554086

**Authors:** Hanna Hõrak, Stephen A. Rolfe, Jurriaan Ton, Julie E. Gray

**Affiliations:** Plants, Photosynthesis & Soil, School of Biosciences, University of Sheffield, Firth Court, Western Bank, Sheffield, S10 2TN, UK; Institute of Technology, University of Tartu, Nooruse 1, 50411, Tartu, Estonia

## Abstract

The phytohormone salicylic acid (SA) is an important molecular signal that mediates pathogen defence mechanisms, including triggering Arabidopsis immune responses to the hemi-biotroph *Pseudomonas syringae* pv. *tomato* (*Pst*). SA induces the expression of a myriad of defence genes via its receptor and transcriptional regulator NONEXPRESSER OF PR GENES 1 (NPR1). Here, we used chlorophyll fluorescence imaging of F_v_/F_m_, to detect damage to photosystem II before *Pst*-induced disease symptoms were visible. We observed that the pathogen only induced damage, and subsequent cell death, in mature leaves while developing leaves in the center of the rosette appeared to be protected. However, in the *npr1-1* mutant, *Pst*-infected mature leaves were able to systemically transmit a signal that caused damage to the photosynthetic machinery in uninfected young leaves. Reductions in F_v_/F_m_ could also be induced systemically in developing *npr1-1* leaves by high levels of SA in mature leaves, and rescued by SA biosynthesis deficiency in *npr1-1sid2-2* mutants. Together, these results indicate that, in addition to its well-known role as a positive regulator of SA responses, NPR1 also acts to suppress SA-dependent immune responses and thereby protects developing leaves from autoimmune damage.

## Introduction

Plant pathogens account for approximately a quarter of major food crop losses globally (Savary et al., 2019). Plant diseases are usually detected by visual inspection, but at the time when disease symptoms become visible to the naked eye, the disease has already progressed. Early disease detection would help in applying appropriate treatment to stop the spread of pathogens. Chlorophyll fluorescence imaging can be applied as a technique to detect plant disease in its early stages, before visual symptoms appear (Pérez-Bueno et al., 2019). The maximum quantum yield of photosystem II (F_v_/F_m_) is a chlorophyll fluorescence parameter that reflects the status of plant photosynthetic machinery (Maxwell and Johnson, 2000). Measurement of F_v_/F_m_ is fast and easy; values are typically ∼0.83 in healthy plants but this is reduced in response to stress (Björkman and Demmig, 1987; Johnson et al., 1993; Maxwell and Johnson, 2000). Fluorescence imaging of pathogen-infected plants can detect a reduction in F_v_/F_m_ before visual disease symptoms occur (Cséfalvay et al., 2009; Kuckenberg et al., 2009; Pérez-Bueno et al., 2016) and F_v_/F_m_ imaging in *Arabidopsis thaliana* (Arabidopsis) has helped to dissect the role of pathogen effectors and abscisic acid (ABA) in plant immunity against the hemi-biotrophic bacterium *Pseudomonas syringae* (de Torres Zabala et al., 2015). Thus, chlorophyll fluorescence imaging is a potent tool for studying plant-pathogen interactions and assessing damage caused by disease at the early stages of infection.

The phytohormone salicylic acid (SA) plays a key role in Arabidopsis immune responses against the bacterium *P. syringae*, and SA levels rise in infected tissues (Summermatter et al., 1994). The majority of SA is produced from chorismate by the isochorismate synthase SALICYLIC ACID INDUCTION DEFICIENT 2 (SID2, Wildermuth et al., 2001). SA is perceived by two types of receptors: NONEXPRESSER OF PR GENES 1 (NPR1, Wu et al., 2012) and NPR3/NPR4 (Fu et al., 2012). NPR1 functions as a major positive regulator of SA responses by activating SA-dependent gene expression via TGACG SEQUENCE-SPECIFIC BINDING PROTEIN (TGA) transcription factors (Zhang et al., 1999; Després et al., 2000; Zhou et al., 2000), whereas repression of SA-dependent gene expression by TGAs and NPR3/NPR4 is released in the presence of SA (Ding et al., 2018).

NPR1 was discovered in a screen for mutants deficient in the induction of systemic acquired resistance (SAR) and expression of *PATHOGENESIS-RELATED* (*PR*) genes (Cao et al., 1994). The *npr1-1* mutant has a point mutation leading to replacement of a conserved histidine at position 334 in the third ankyrin-repeat of the protein by a tyrosine residue (Cao et al., 1997). While *npr1-1* is not a null allele and a mutant version of the protein is expressed in these plants (Ding et al., 2020), interaction with TGA transcription factors (Zhang et al., 1999; Després et al., 2000; Zhou et al., 2000) and SA-inducible *PR* gene expression are severely repressed in *npr1-1* (Cao et al., 1994), leading to enhanced susceptibility to *P. syringae* infection.

Although NPR1 has been considered a major positive regulator of SA-responses for many years, there is evidence that NPR1 may also contribute to suppressing SA biosynthesis and SA-dependent signaling. For example, *npr1* mutants contain higher levels of SA both in the unchallenged state (Delaney et al., 1995; Clarke et al., 2000) and after pathogen infection (Delaney et al., 1995; Wildermuth et al., 2001; Zhang et al., 2010; DeFraia et al., 2013; Liu et al., 2020) or ozone treatment (Ogawa et al., 2007). NPR1 also suppresses systemic changes in circadian clock rhythms caused by local bacterial infection or SA treatment, as *npr1-1* mutants were hypersensitive to respective treatments (Li et al., 2018). Thus, in addition to activating SA-dependent defense gene expression, NPR1 may also have a negative regulatory role in SA signaling.

Here we used chlorophyll fluorescence imaging to characterize local and systemic effects of *Pst* infection on the F_v_/F_m_ values of *npr1-1* and wild-type plants. We show that in addition to being a well-known positive regulator of SA responses, NPR1 is also required for suppressing SA-dependent immune responses in immature leaves. We propose that NPR1 functions in this way to protect the photosynthetic machinery of developing leaves from damage triggered by systemic induction of immune responses by high levels of SA accumulation in mature leaves.

## Results

### Chlorophyll fluorescence imaging detects disease before visual symptoms appear

First, we confirmed that, under our conditions, whole plant chlorophyll fluorescence imaging could detect damage caused by the bacterial pathogen *Pseudomonas syringae* pv. *tomato* strain DC3000 (*Pst*) before signs of cell death caused by hypersensitive response became visible in Arabidopsis leaves. Dip-inoculation of wild-type Arabidopsis with *Pst* led to a clear decrease in maximum quantum yield of photosystem II (F_v_/F_m_) in mature leaves. At 24 hours post inoculation (hpi), F_v_/F_m_ had dropped to ∼0.73 in mature leaves of *Pst*-infected plants, compared with ∼0.83 in mock-treated plants indicating that damage had occurred, although the plants remained indistinguishable to the naked eye (Figure 1A). Three days after inoculation, the areas that had reduced F_v_/F_m_ at 24 hpi had developed visible lesions (Figure 1A), indicating that chlorophyll fluorescence imaging had enabled us to detect early bacterial infection-induced damage before visual symptoms appeared.

**Figure 1.**
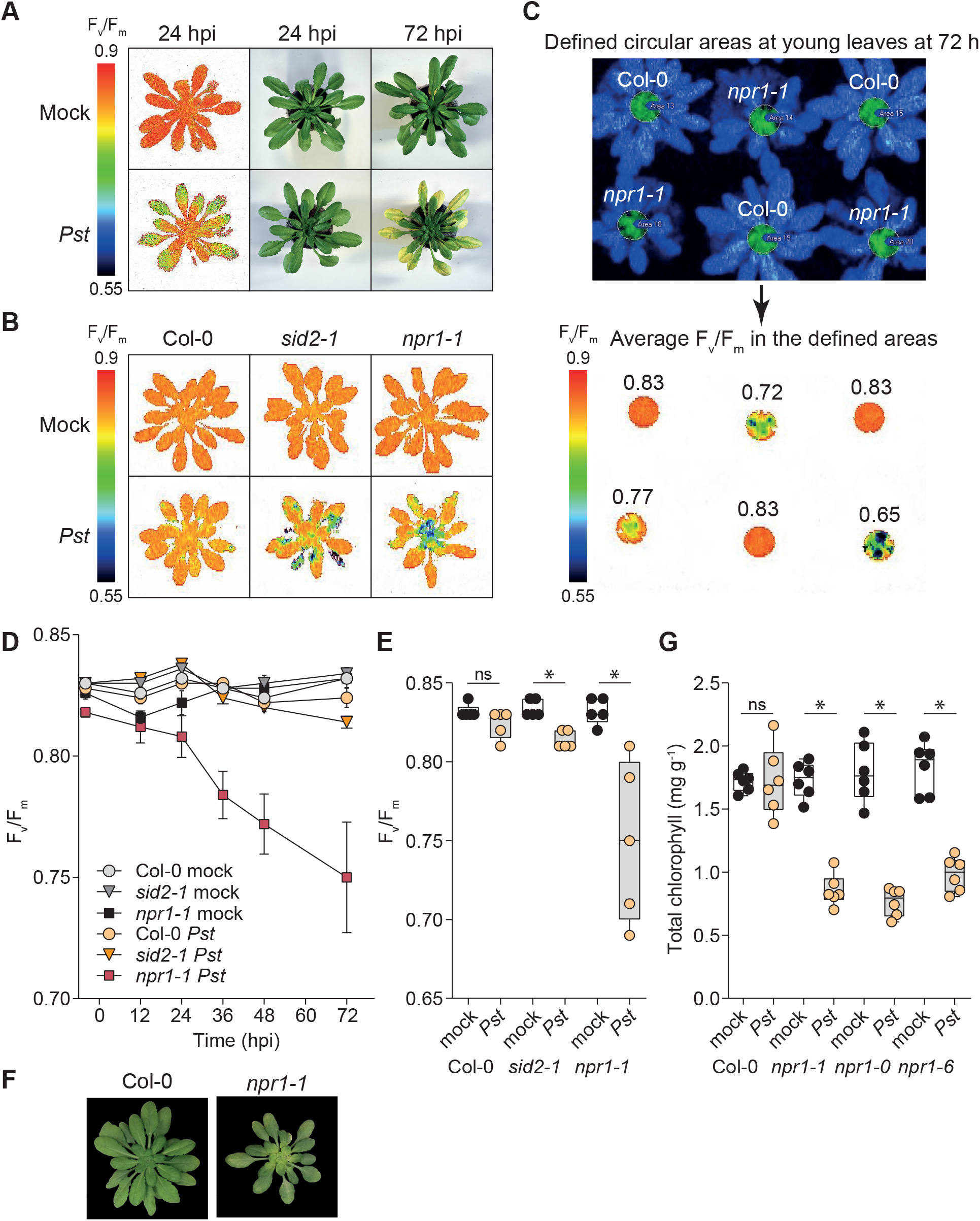
*P. syringae* infection induces reduction of F_v_/F_m_ in developing leaves of *npr1-1*. **(A)** Representative F_v_/F_m_ images and photos of wild-type Arabidopsis dip-inoculated with *Pst* or mock-inoculated 24 and 72 hours post inoculation (hpi). **(B)** Representative F_v_/F_m_ images of plants spray-inoculated with *Pst* at 72 hpi. **(C)** Illustration of quantification of F_v_/F_m_ in developing leaves. **(D)** Time course of F_v_/F_m_ in developing leaves from 6 h before to 72 h after spray-inoculation with *Pst* and **(E)** F_v_/F_m_ in developing leaves at 72 hpi, n = 5 plants. **(F)** Representative images of plants spray-inoculated with *Pst* 5 days after inoculation. **(G)** Chlorophyll content of developing leaves 5 days after spray inoculation, n = 6 plants. In **(E)** and **(G)**, dots show measurements from individual plants and stars show statistically significant differences between groups (Welch’s t-test with Bonferroni correction).

### Bacterial infection leads to reduction of F_v_/F_m_ in young leaves of npr1-1

We noticed that the *Pst*-induced decrease in F_v_/F_m_ and subsequent cell death were present only in mature leaves, and not in young developing leaves at the center of the Arabidopsis rosette (Figure 1A). Younger Arabidopsis leaves are known to accumulate higher levels of SA and activate SAR more effectively than older leaves during *P. syringae* infection (Zeier, 2005). As SA has a key role in defense responses triggered by *Pst*, we tested whether SA biosynthesis or signaling through NPR1 are required for the protection of young leaves from damage caused by *Pst* infection. We analyzed changes in F_v_/F_m_ over three days after spray-inoculation with *Pst* in the *sid2-1* and *npr1-1* mutants that are deficient in SA biosynthesis or SA-dependent gene expression induction via NPR1, respectively. Under these conditions, *Pst* did not cause strong disease symptoms in wild-type plants, but triggered a reduction of F_v_/F_m_ in mature and to a smaller extent also in developing leaves of *sid2-1* mutants (Figure 1B), suggesting that SA-deficiency results in increased susceptibility to infection in all leaves of SA-deficient plants. In contrast, we detected a clear reduction of F_v_/F_m_ in response to *Pst* inoculation in *npr1-1* that was most notable in the young leaves at the center of the rosette (Figure 1B), indicating that a functional NPR1 is required to protect young leaves from bacterial infection-induced damage. We monitored F_v_/F_m_ of young leaves in a circular area at the center of Arabidopsis rosettes (Figure 1C) and found that F_v_/F_m_ decreased from ∼0.83 to ∼0.75 over three days following spray-inoculation with *Pst* in the *npr1-1* mutants (Figure 1D, E). Five days after inoculation with *Pst*, the damage indicated by previous reduction of F_v_/F_m_ showed in the increasingly pale green color of the young leaves of *npr1-1* (Figure 1F), suggesting reduced chlorophyll levels. To confirm that the phenotype is due to NPR1 loss-of-function, we additionally analyzed two T-DNA insertion lines deficient in NPR1: *npr1-0* and *npr1-6*. We quantified total chlorophyll levels in the young leaves at the center of Arabidopsis rosettes 5 days after spray-inoculation with *Pst* and found a significant decrease in chlorophyll content in all *npr1* mutants studied in comparison to wild-type (Figure 1G). Thus, NPR1 has a role in protecting young leaves from *Pst* infection-induced damage.

### Bacterial infection in mature leaves triggers systemic reduction of F_v_/F_m_ in young leaves of npr1-1

We next asked, whether local bacterial infection in mature leaves is sufficient to trigger reduction of F_v_/F_m_ in the young leaves of *npr1-1* via a systemic signaling mechanism. We infiltrated three mature leaves of plants with *Pst* at OD_600_=0.02 or OD_600_=0.2 and monitored F_v_/F_m_ in young leaves for three days. In *npr1-1*, infection of mature leaves with *Pst* at OD_600_=0.02 caused a clear decrease in F_v_/F_m_ in developing leaves that was not present in plants that were mock-treated or infiltrated with a higher-density inoculum (Figure 2A, B and C). These data suggest that bacterial infection in mature leaves triggers a systemic response in the young leaves that leads to a reduction in F_v_/F_m_ in the *npr1-1* mutant that was not seen in the wild-type plants. This systemic response did not occur in the *npr1-1* plants that were treated with higher numbers of bacteria, suggesting that the generation and movement of the systemic signal from infected tissue requires time and may not occur in heavily infected leaves that die quickly.

**Figure 2.**
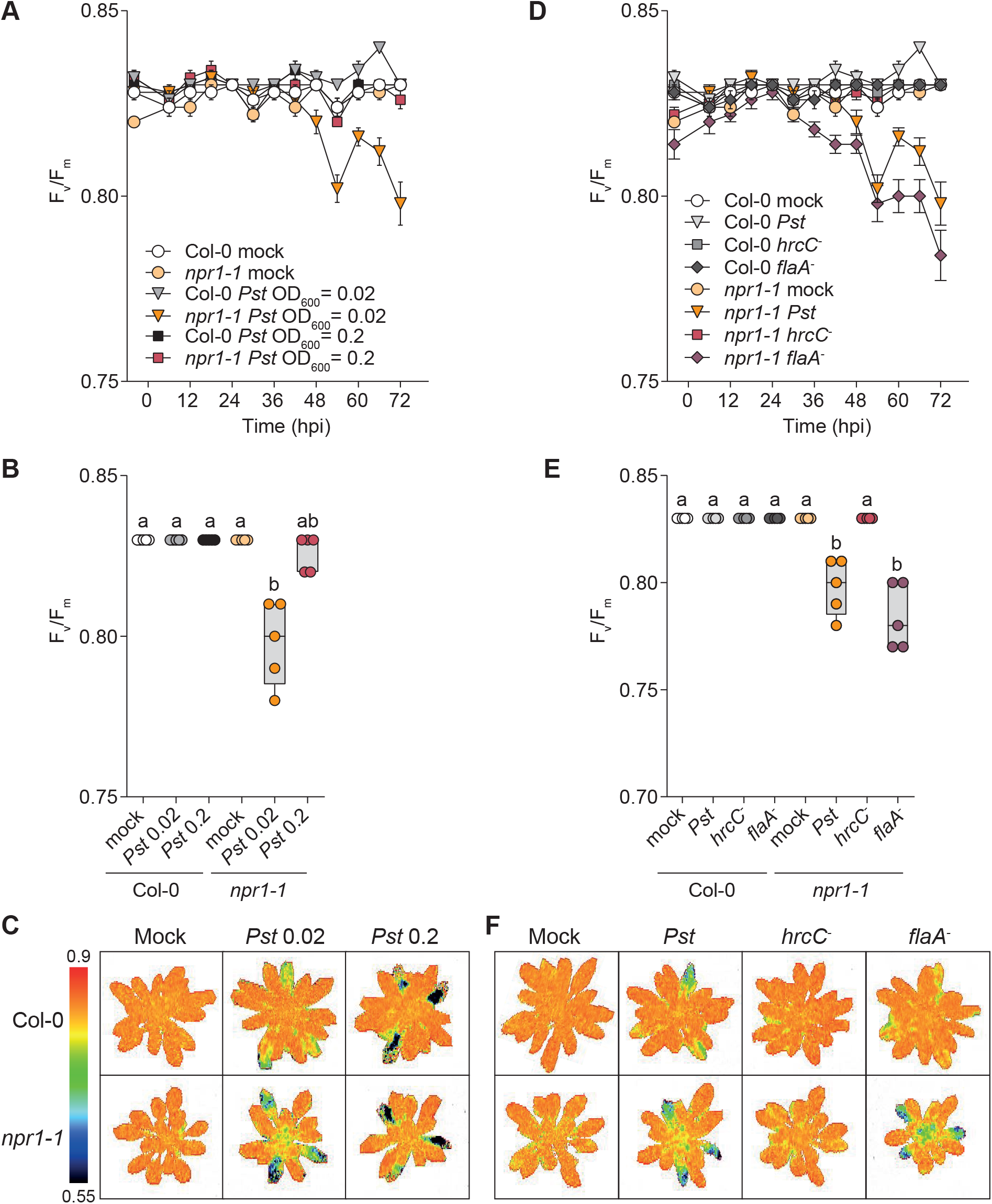
Local effector-producing *P. syringae* infection in mature leaves induces reduction of F_v_/F_m_ in developing leaves of *npr1-1*. **(A)** Time course of F_v_/F_m_ in developing leaves from 6 h before to 72 h after infiltration of three mature leaves with *Pst* at indicated inoculum densities and **(B)** F_v_/F_m_ in developing leaves at 72 hpi, mean ± SEM is shown, n = 5 plants. **(C)** Representative F_v_/F_m_ images of plants 72 hours after three mature leaves were inoculated with *Pst* at indicated inoculum densities. **(D)** Time course of F_v_/F_m_ in developing leaves from 6 h before to 72 h after infiltration of three mature leaves with indicated *Pst* strains at OD_600_=0.02 and **(E)** F_v_/F_m_ in developing leaves at 72 hpi, mean ± SEM is shown, n = 5 plants. **(F)** Representative F_v_/F_m_ images of plants 72 hours after three mature leaves were inoculated with indicated *Pst* strains at OD_600_=0.02. In **(B)** and **(E)** dots show measurements from individual plants and letters show statistically significant differences between groups (Kruskal-Wallis with Dunn *post hoc* test).

To test whether the observed systemic reduction of F_v_/F_m_ in the developing leaves of the *npr1-1* mutant was dependent on flagellin-triggered responses, we inoculated three mature leaves of wild-type and *npr1-1* plants with the flagellin-deficient *flaA*^*-*^ strain (Hu et al., 2001) of *Pst* at OD_600_=0.02. Monitoring of F_v_/F_m_ in developing leaves over three days showed that the *Pst flaA*^*-*^ strain triggered a systemic reduction in F_v_/F_m_ in *npr1-1* but not wild-type plants, similar to *Pst* (Figure 2D, E and F). These results indicate that flagellin-dependent signaling is not required for the systemic response that leads to the reduction in F_v_/F_m_ in the *npr1-1* mutant. We also tested whether bacterial effectors are required to trigger the systemic reduction of F_v_/F_m_ in young leaves of the *npr1-1* mutant. To this end, we inoculated three mature leaves of *npr1-1* or wild-type plants with the *hrcC*^*-*^ strain (Yuan and He, 1996) of *Pst* that cannot deliver effectors to plant cells due to lack of a functional type III secretion system. There was no change in F_v_/F_m_ in the developing leaves of *Pst hrcC*^*-*^-inoculated *npr1-1* plants during three days after inoculation (Figure 2D, E and F), indicating that bacterial effectors are required to trigger a systemic reduction of F_v_/F_m_ in the developing leaves of *npr1-1*.

### High SA levels in mature leaves trigger systemic reduction of F_v_/F_m_ in npr1-1

*Pst* infection leads to SA accumulation at the infection site (Summermatter et al., 1994), and effector-producing bacteria induce stronger infection and higher levels of SA (Hamdoun et al., 2013). As *npr1-1* is hypersensitive to SA (Cao et al., 1997; Kinkema et al., 2000), we hypothesized that high levels of SA that accumulate in mature leaves in response to bacterial infection may lead to the observed reduction of F_v_/F_m_ in the developing leaves of *npr1-1*. To test this hypothesis, we infiltrated three mature leaves of *npr1-1* and wild-type plants with SA at 0.5 mM or 1 mM concentration and monitored F_v_/F_m_ in developing leaves. 1 mM SA triggered a significant reduction in F_v_/F_m_ that was evident during 54 hours after infiltration (Figure 3A, B and C) in developing leaves of *npr1-1*, but not wild-type plants. Thus, high levels of SA in mature leaves are sufficient to systemically trigger a reduction of F_v_/F_m_ in the young developing leaves of *npr1-1*.

**Figure 3.**
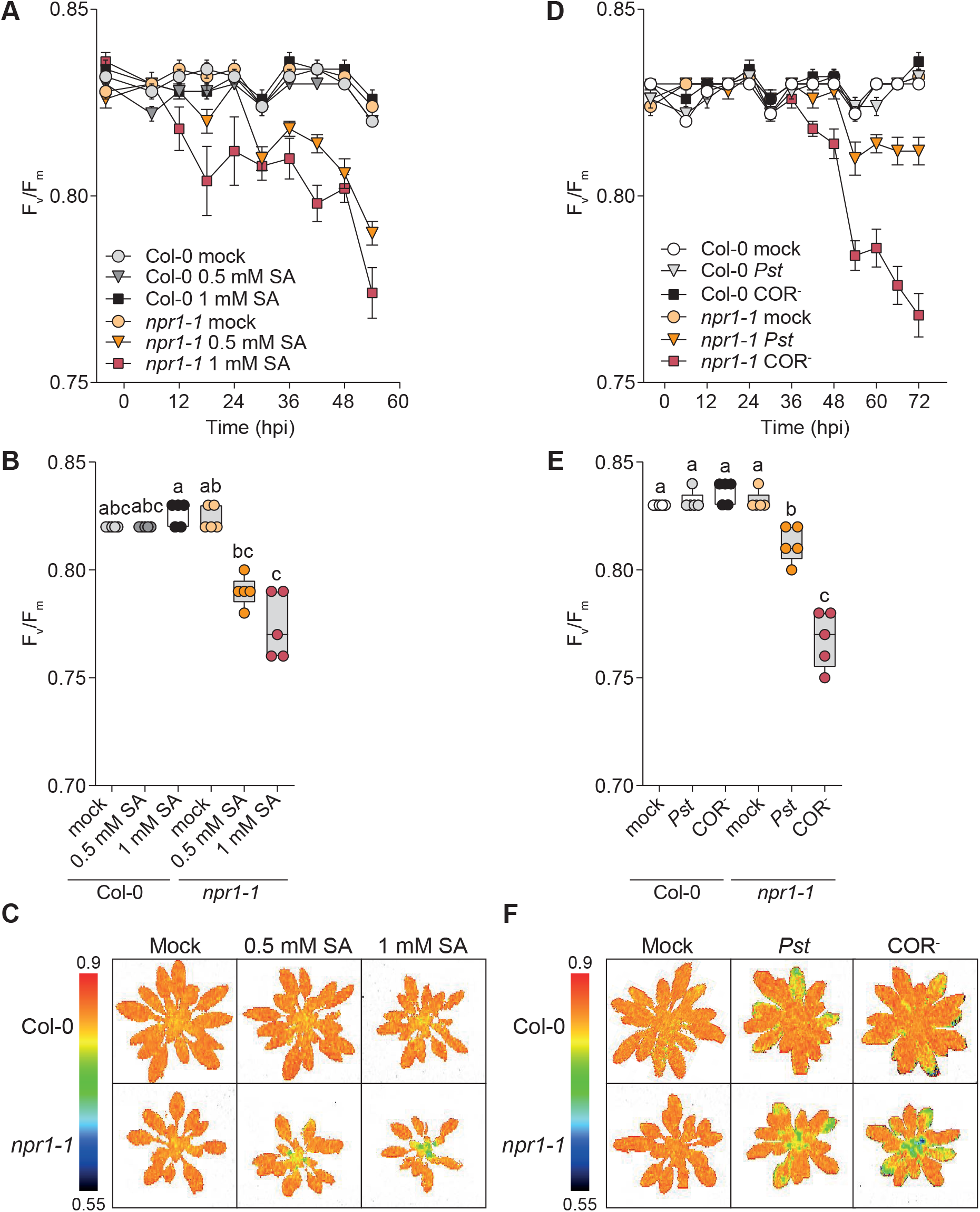
High SA levels in mature leaves induce reduction of F_v_/F_m_ in developing leaves of *npr1-1*. **(A)** Time course of F_v_/F_m_ in developing leaves from 6 h before to 54 h after infiltration of three mature leaves with SA at indicated concentrations and **(B)** F_v_/F_m_ in developing leaves at 54 hpi, mean ± SEM is shown, n = 5 plants. **(C)** Representative F_v_/F_m_ images of plants 54 hours after three mature leaves were infiltrated with SA at indicated concentrations. **(D)** Time course of F_v_/F_m_ in developing leaves from 6 h before to 72 h after infiltration of three mature leaves with *Pst* or the coronatine-deficient (COR^-^) *Pst* strain at OD_600_=0.02 and **(E)** F_v_/F_m_ in developing leaves at 72 hpi, mean ± SEM is shown, n = 5 plants. **(F)** Representative F_v_/F_m_ images of plants 72 hours after three mature leaves were inoculated with indicated *Pst* strains at OD_600_=0.02. In **(B)** and **(E)** dots show measurements from individual plants and letters show statistically significant differences between groups (Kruskal-Wallis with Dunn *post hoc* test for **(B)** and two-way ANOVA with Tukey *post hoc* test for **(E)**).

To further assess the role of SA in the systemic reduction of F_v_/F_m_ in developing leaves of *npr1-1*, we inoculated three mature leaves of *npr1-1* or wild-type plants with the coronatine deficient COR^-^ *Pst* strain (Brooks et al., 2004). The bacterial effector coronatine mimics jasmonate-isoleucine (Melotto et al., 2008), thus activating jasmonate-dependent defense responses that suppress SA levels (Zheng et al., 2012). We hypothesized that infection caused by the COR^-^ *Pst* strain would prevent the suppression of SA responses via the coronatine-induced JA-pathway, leading to higher SA levels and stronger reduction of F_v_/F_m_ in the young leaves of *npr1-1*. Indeed, *Pst* COR^-^ infection caused a clearly stronger reduction of F_v_/F_m_ in the developing leaves of *npr1-1* than coronatine-producing *Pst* during three days after inoculation of mature leaves (Figure 3D, E and F). Thus, higher accumulation of SA during infection in mature leaves enhances the systemic reduction of F_v_/F_m_ in developing leaves of the *npr1-1* mutant.

### SA biosynthesis deficiency rescues bacteria-induced systemic reduction of F_v_/F_m_ in npr1-1sid2-2

The reduction of F_v_/F_m_ in the developing leaves of *npr1-1* that we detected via chlorophyll fluorescence analysis was a forerunner of chlorosis that became visually detectable ∼5 days after inoculation of plants with bacteria (Figure 1F). Similar chlorosis has been reported before in young leaves of plants carrying the *npr1-1* allele together with mutations leading to high SA levels but not in mutants combining the *npr1-1* allele with low SA levels (Zhang et al., 2008). We hypothesized that reduction of SA levels via impairment of SA biosynthesis in the *npr1-1* background would rescue the phenotype of *npr1-1*. To test this, we analyzed *Pst*-induced changes in chlorophyll fluorescence in the *npr1-1sid2-2* double mutant that is deficient in SA biosynthesis. The systemic reduction of F_v_/F_m_ in developing leaves of *npr1-1* mutants triggered by inoculation of three mature leaves with *Pst* was not present in the SA-biosynthesis deficient *npr1-1sid2-2* mutant (Figure 4A, B and C). The visually detectable chlorosis that was present in young leaves of *npr1-1* plants 5 days after *Pst* inoculation was also absent in the *npr1-1sid2-2* double mutant (Figure 4D). These data further indicate that high levels of SA in the mature leaves of *npr1-1* mutant trigger systemic damage of photosynthetic machinery in the developing leaves and suggest that in wild-type plants, NPR1 acts as a negative regulator of SA levels and/or responses, protecting developing leaves from damage caused by excessive defense responses (Figure 5).

**Figure 4.**
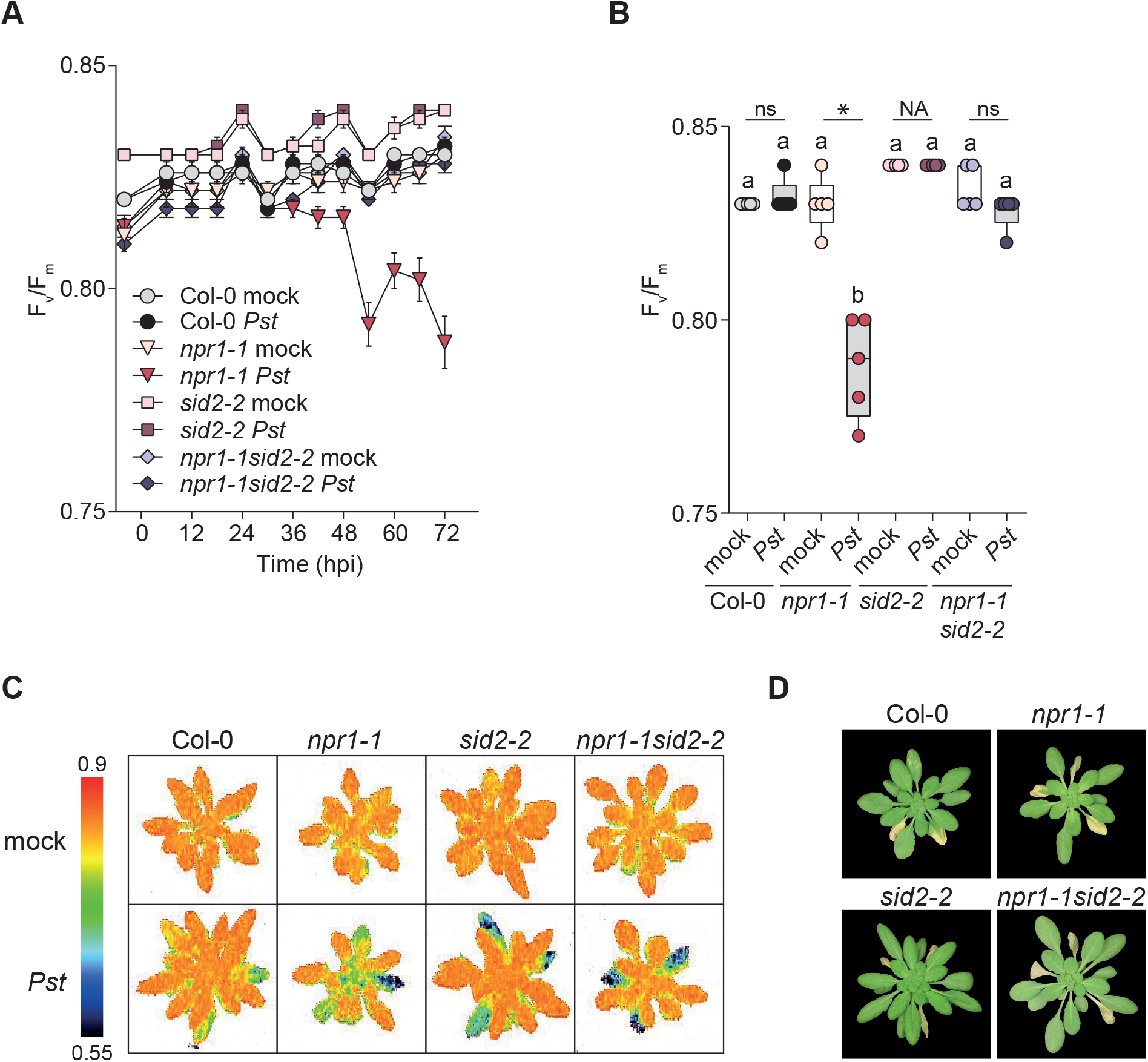
SA biosynthesis deficiency rescues bacteria-induced systemic reduction of F_v_/F_m_ in *npr1-1sid2-2* double mutant. **(A)** Time course of F_v_/F_m_ in developing leaves from 6 h before to 72 h after infiltration of three mature leaves with *Pst* at OD_600_=0.02 and **(B)** F_v_/F_m_ in developing leaves at 72 hpi, mean ± SEM is shown, n = 5 plants. **(C)** Representative F_v_/F_m_ images of plants 72 hours after three mature leaves were infiltrated with *Pst* at OD_600_=0.02. **(D)** Representative images of plants 5 days after three mature leaves were inoculated with *Pst* at OD_600_=0.02. In **(B)**, dots show measurements from individual plants and stars show statistically significant differences between groups (Welch’s t-test with Bonferroni correction).

**Figure 5.**
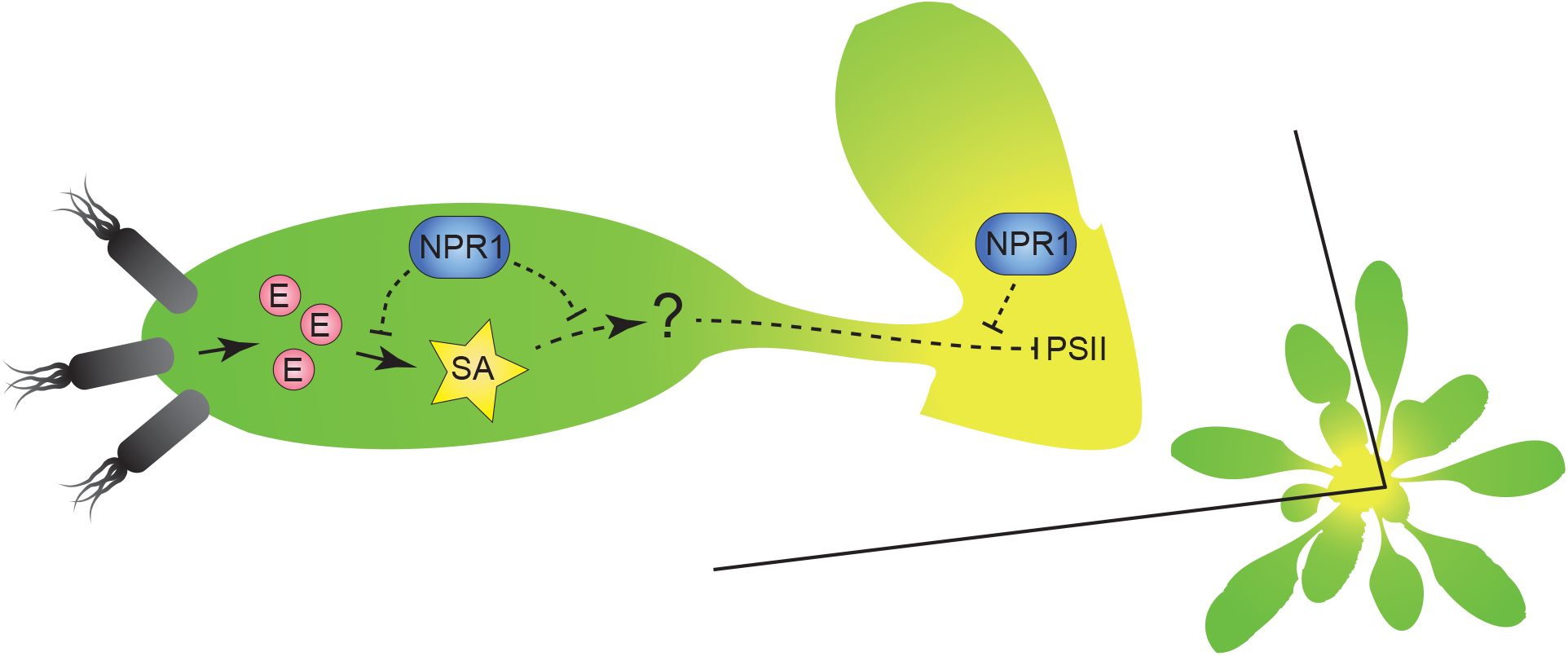
Model for a negative regulatory role of NPR1 in SA-induced responses during *Pst* infection. Bacterial infection induces production of SA in infected tissues, whereas virulent effector-producing (E) strains lead to higher SA levels. High levels of SA in mature leaves lead to induction of systemic signals (SA itself or unknown signals indicated by the question mark) that are suppressed by NPR1 via suppression of SA production and/or propagation or amplification of the signal. NPR1 could act at one or all of the points indicated in the proposed pathway. In the *npr1-1* mutant the negative regulator is dysfunctional, ultimately leading to autoimmune damage to photosystem II (PSII) in developing leaves manifested initially as reduction of F_v_/F_m_ and later as chlorosis.

## Discussion

We used chlorophyll fluorescence imaging to detect *P. syringae*-induced damage in Arabidopsis and detected a reduction in F_v_/F_m_ in infected leaves before visual disease symptoms (Figure 1A), indicating bacterial-induced damage to PSII. In addition to symptoms caused by bacteria in infected tissues, we also detected systemic infection-induced damage to the photosynthetic machinery in uninfected developing leaves of the *npr1-1* mutant (Figures 2, 3, 4). The systemic damage was triggered by high SA levels in mature leaves generated during infection (Figure 3) and was rescued in the SA biosynthesis-deficient *npr1-1sid2-2* mutant (Figure 4). These findings suggest a negative regulatory role for NPR1 in protecting developing leaves from damage caused by excessive activation of SA-dependent immune responses (Figure 5).

When *npr1-1* seedlings are grown on medium with SA they bleach and die (Cao et al., 1997; Kinkema et al., 2000), indicating that the *npr1-1* mutant is hypersensitive to SA. Several mutants that carry the *npr1-1* allele together with alleles resulting in high SA levels display chlorosis in young leaves (Zhang et al., 2008), similar to the systemic *Pst*-induced damage found here, and supporting our hypothesis that developing leaves of *npr1-1* are especially sensitive to SA. High SA levels in mature leaves are sufficient to cause a reduction in F_v_/F_m_ in young leaves of *npr1-1* (Figure 3), indicating that SA either serves as, or triggers, a systemic signal that results in damage to photosystem II in plants where NPR1 function is impaired. As *npr1-1* is not a null allele, but activation of SA-dependent gene expression via TGA transcription factors is impaired in this mutant, the phenotypes observed here suggest that NPR1-dependent regulation of gene expression is responsible for preventing SA-induced damage to photosynthetic machinery in developing leaves. In line with this, nuclear localization of NPR1 is required to prevent SA-hypersensitivity in seedlings (Zhang et al., 2010) and the seedlings of TGA transcription factor triple mutant *tga2tga5tga6* are also more sensitive to SA (Zhang et al., 2003).

Our experiments raise the question, what is the systemic signal that ultimately leads to damage to PSII and chlorosis in *npr1-1* plants with high SA levels in mature leaves – is SA itself the mobile signal or is it a different molecule? Our data suggest that the generation and propagation of this systemic signal requires time, as *npr1-1* plants infected with a high inoculum of *Pst* that killed mature leaves quickly did not develop damage to PSII in young leaves (Figure 2A-C). Possibly, a threshold of SA needs to be reached in mature leaves, before either another systemic signal is triggered, or so that SA levels are sufficient for movement to distal tissues and elicitation of a systemic response. During SAR, SA and the mobile signal *N*-hydroxy-pipecolic acid (NHP) accumulate in local tissues in response to infection, and both the ability to produce SA and movement of NHP to systemic tissues are needed for SAR induction (Nawrath and Métraux, 1999; Chen et al., 2018; Hartmann et al., 2018; Mohnike et al., 2021). As pathogen-induced NHP levels are higher in *npr1-1* than in wild-type plants (Hartmann et al., 2018; Liu et al., 2020), a potential role for NHP in SA-hypersensitivity of *npr1-1* merits further study, for example via analysis of *npr1-1* plants deficient in NHP biosynthesis. The enzyme UGT76B1 glycosylates both SA and NHP (Noutoshi et al., 2012; Bauer et al., 2021). Pathogen- and SA-induced expression of *UGT76B1* transcript was lower in *npr1-1* mutants compared with wild-type plants (Liu et al., 2020). Thus, it is possible that the inability to efficiently convert SA and/or NHP into their glycosylated forms that lack immune-stimulating activity is responsible for pathogen- and SA-induced damage in *npr1* mutants.

The systemic SA-induced damage to PSII that later manifested as chlorosis in *npr1-1* resembles meristem chlorosis observed in plants that either express the *P. syringae* effector protein HopAM1 or have been infected with bacteria producing this effector (Goel et al., 2008). Similarly to the SA-induced reduction of F_v_/F_m_ in the developing leaves of *npr1-1*, meristem chlorosis in Arabidopsis infected with HopAM1-producing bacteria is also believed to be systemically transmitted, as no bacteria were recovered from the developing chlorotic leaves (Goel et al., 2008). The HopAM1-induced meristem chlorosis was absent in the *rbohdrbohf* double mutant, indicating that reactive oxygen species (ROS) production via apoplastic NADPH oxidases contributes to systemic chlorosis (Iakovidis et al., 2016). Likewise, it is possible that higher SA-dependent induction of ROS leads to damage to PSII in developing *npr1-1* leaves. Overexpression of FLAVIN-DEPENDENT MONOOXYGENASE 1 (FMO1) – the enzyme that catalyzes the biosynthesis of NHP from pipecolic acid – leads to higher ROS accumulation (Chen and Umeda, 2015). Thus, high NHP levels in *npr1-1* (Hartmann et al., 2018; Liu et al., 2020) may contribute towards damage to PSII though ROS in the developing leaves of this mutant. Further experiments addressing ROS levels and using ROS-production deficient *npr1-1* mutants will help to clarify the role of ROS in systemic SA-induced chlorosis in *npr1-1*.

SA or *Pst* infection of *npr1-1* mutants is known to trigger enhanced systemic changes in SA-mediated circadian clock rhythms, suggesting a negative regulatory role for NPR1 in control of SA responses (Li et al., 2018). The systemic autoimmune damage that occurs in young leaves of *npr1-1* mutants reported here further demonstrates that NPR1 acts as a suppressor of systemic SA-mediated responses, in addition to its better-known role as the major positive regulator of SAR. These findings underline the complexity of SA signaling and the necessity for research into this negative regulatory role of the NPR1 signaling hub in responses mediated by SA.

Chlorophyll fluorescence imaging has been previously applied to detect pathogen-induced damage in various plant species. Our experiments show that this technique can be used for early detection of damage to photosynthesis caused by disease, but also by excessive immune response in Arabidopsis. Here we only measured the fast and simple chlorophyll fluorescence parameter F_v_/F_m_ but measurement and calculation of more complex parameters could perhaps give further temporal and spatial insight into disease progression and immune responses during the *P. syringae*-Arabidopsis interaction.

## Materials and methods

### Plant material and growth conditions

*Arabidopsis thaliana* (Arabidopsis) Col-0 accession and the following mutants in the same genetic background were used in experiments: *npr1-0* (SALK_204100 (Alonso et al., 2003; Skelly et al., 2019)), *npr1-1* (Cao et al., 1994), *npr1-6* (SAIL_708_F09, (Sessions et al., 2002; Xin et al., 2016)), *sid2-1* (Nawrath and Métraux, 1999), *sid2-2* (Dewdney et al., 2000), *npr1-1sid2-2* (Nair et al., 2021). Plants were grown in 3:1 v/v mixture of compost (Levington M3) and perlite in a controlled environment chamber (Conviron) with 9/15 h day/night regime (22°C day, 18°C night), 200 μmol m^-2^ s^-1^ of light and 60% relative air humidity (RH), 5-week-old plants were used for experiments.

### Bacterial strains, growth conditions, pathogen and chemical treatments

*Pseudomonas syringae* pv. *tomato* DC3000 (*Pst* DC3000) and its coronatine-deficient (Brooks et al., 2004), effector-secretion deficient *hrcC* (Yuan and He, 1996) and flagellin-deficient *fliC* (*flaA*) (Hu et al., 2001) strains were grown in King’s B medium at room temperature with 50 μg ml^-1^ Rifampicin overnight to an OD_600_ of 0.15 to 0.8. Bacteria were collected by centrifugation and re-suspended in 10 mM MgSO_4_ to an OD_600_ of 0.2 (spray or dip inoculation) or 0.02 (syringe infiltration). For spray or dip inoculation, Silwet-L77 (De Sangosse, http://www.desangosse.com) was added to bacterial suspension at the final concentration of 0.02% and control (mock-treated) plants were treated with 0.02% Silwet-L77 in 10 mM MgSO_4_. For syringe infiltration, 10 mM MgSO_4_ was the mock treatment. For salicylic acid treatment, 0.5 mM or 1 mM sodium salicylate (Sigma, https://www.sigmaaldrich.com) with 0.01% ethanol was infiltrated into mature leaves, 0.01% ethanol solution was used as mock treatment.

### Chlorophyll fluorescence analysis

Chlorophyll fluorescence measurements were carried out with a PlantScreen chlorophyll fluorescence scanner (Photon Systems Instruments, https://psi.cz/) set up inside the controlled environment chamber where plants were grown and treated. Dark-adapted F_v_/F_m_ was measured before treatment and with 6h intervals after treatment using the manufacturer’s protocol with the following settings: excitation light Act1 at 102 μmol m^-2^ s^-1^ and Super at 909 μmol m^-2^ s^-1^ (40% and 25% of respective maximal values), 20% sensitivity and 20 μs shutter speed. Average F_v_/F_m_ values for young leaves were obtained by defining appropriate regions of interest at rosette centers and extracting respective F_v_/F_m_ values with the FluorCam7 software (Photon Systems Instruments, https://psi.cz/).

### Chlorophyll content measurement

Plants for chlorophyll content measurement were grown in 2:1 v/v mixture of peat (Kekkilä) and vermiculite in a controlled growth room with 10/16 h day/night regime (23°C day, 19°C night), 250 μmol m^-2^ s^-1^ of light and 60% RH. Five days after spray inoculation with *Pst*, young leaves at the center of the rosette were excised, weighed and ground in liquid nitrogen. Chlorophyll was extracted with acetone and total chlorophyll content determined as in Lichtenthaler and Buschmann (2001).

### Statistical analysis

Kruskal-Wallis with Dunn *post hoc* test, two-way ANOVA with Tukey *post-hoc* test or Welch’s t-test with Bonferroni correction for multiple testing were used as appropriate and as indicated in the figure legends. Effects were considered significant at p < 0.05, or using the Bonferroni-corrected significance levels in Fig. 1E, 1G and 4B.

## Acknowledgments

We thank Christiane Gatz for *npr1-1sid2-2* seeds and Cyril Zipfel for sharing the *flaA*^*-*^, *hrcC*^*-*^ and COR^-^ *P. syringae* strains. This work was supported by Leverhulme Trust grant RPG-2016-274 to J.E.G. and the Estonian Research Council grant PSG404 to H.H. We are grateful for funding from the Leverhulme Trust and Wolfson Foundation.

## Author Contributions

H.H designed and performed experiments and analyzed data with advice from S.A.R, J.T and J.E.G; H.H wrote the manuscript, all authors commented, edited and approved the final manuscript.

## Notes

### Competing Interest Statement

The authors have declared no competing interest.

